# OpenABC Enables Flexible, Simplified, and Efficient GPU Accelerated Simulations of Biomolecular Condensates

**DOI:** 10.1101/2023.04.19.537533

**Authors:** Shuming Liu, Cong Wang, Andrew Latham, Xinqiang Ding, Bin Zhang

## Abstract

Biomolecular condensates are important structures in various cellular processes but are challenging to study using traditional experimental techniques. In silico simulations with residue-level coarse-grained models strike a balance between computational efficiency and chemical accuracy. They could offer valuable insights by connecting the emergent properties of these complex systems with molecular sequences. However, existing coarse-grained models often lack easy-to-follow tutorials and are implemented in software that is not optimal for condensate simulations. To address these issues, we introduce OpenABC, a software package that greatly simplifies the setup and execution of coarse-grained condensate simulations with multiple force fields using Python scripting. OpenABC seamlessly integrates with the OpenMM molecular dynamics engine, enabling efficient simulations with performances on a single GPU that rival the speed achieved by hundreds of CPUs. We also provide tools that convert coarse-grained configurations to all-atom structures for atomistic simulations. We anticipate that OpenABC will significantly facilitate the adoption of in silico simulations by a broader community to investigate the structural and dynamical properties of condensates. Open-ABC is available at https://github.com/ZhangGroup-MITChemistry/OpenABC

## Introduction

Biomolecular condensates underly the organization of many cellular processes, such as speckles for RNA splicing, nucleoli for ribosomal RNA processes, P granule for stress response, etc.^1–14^ They are also termed membrane-less organelles due to the lack of enclosure and exhibit liquid-like properties. Intrinsically disordered proteins (IDPs) and RNA molecules are enriched inside the condensates.^3,4,9,11^ These molecules promote promiscuous, multivalent interactions, leading to spontaneous phase transition and condensate formation.^15^ The nature of the molecular interactions that drive phase separation, the microenvironment of the condensates, and their dynamical relaxation, are under active investigation.

Computational modeling can prove invaluable for studying biomolecular condensates by providing detailed structural and dynamic characterizations.^16–42^ Particle-based coarsegrained modeling approaches are promising since their computational efficiency enables long timescale simulations to promote large-scale reorganization for structural relaxation.^43–46^ Such simulations may predict condensate physical properties de novo, elucidating the connection between molecular sequences and emergent properties.^18,47^ However, the reduced resolution of these coarse-grained models could be insufficient for describing the complex microenvironment of condensate interior.^48–50^ Atomistic simulations with explicit representation of solvent molecules and counter ions can be necessary for further characterizing physicochemical interactions that produce the selective partition of small molecules within condensates.^49–53^ Combining the two modeling approaches at different resolutions could be particularly powerful since they enable long-timescale simulations for structural relaxation while preserving the fine-resolution details.

While many computational models and force fields have been introduced for simulations of IDPs and biomolecules, software engineering has yet to catch up. There is an urgent need to build user-friendly tools to set up and execute condensate simulations. Preparing biomolecular simulations can be rather involved. Even creating initial configurations for such simulations is often non-trivial. Much-dedicated software has been introduced to prepare atomistic simulations,^54–57^ and existing molecular dynamics (MD) simulation packages are highly optimized for computational efficiency. ^54,57–59^ However, existing tools are not immediately transferable for setting up coarse-grained condensate simulations. Furthermore, coarse-grained force fields are often implemented into disparate simulation engines not necessarily best suited for condensate simulations, hindering cross-validation and the unleashing of full modeling potential. Further software development can significantly reduce the entry barrier for in silico studies, allowing more researchers to experience the usefulness of computational modeling. They could facilitate comparing and benchmarking various force fields, driving continuous improvement.

We introduce a software package termed OpenABC for “**Open**MM GPU-**A**ccelerated simulations of **B**iomolecular **C**ondensates”. The package is flexible and implements multiple popular coarse-grained force fields for simulations, including the hydropathy scale (HPS) protein model, the maximum entropy optimized force field (MOFF) for proteins, and the molecular renormalization group (MRG)-CG DNA model.^20,28,60,61^ It dramatically simplifies the simulation setup: only a few lines of Python scripts are needed to carry out condensate simulations starting from initial configurations of a single protein or DNA. The package is integrated with OpenMM, a GPU-accelerated MD engine,^62^ enabling efficient simulations with advanced sampling techniques. Finally, we include tools that allow converting coarsegrained configurations to atomistic structures for further condensate modeling with all-atom force fields. Tutorials in Jupyter Notebooks are provided to demonstrate the various capabilities. We anticipate OpenABC to greatly facilitate the application of existing computer models for simulating biomolecular condensates and the continued force field development.

## Methods

### Details of molecular dynamics simulations

We performed temperature replica-exchange simulations^63^ with MOFF to determine the conformational ensembles of HP1*α* and HP1*β* dimers. Atomistic protein structures were predicted with RaptorX^64^ and used to initialize simulations. Details on modeling HP1 proteins to preserve the tertiary structure of folded domains are provided in the Supporting Information *Section: Setting up MOFF HP1 system*. Six independent replicas were simulated to maintain temperatures at 300 K, 315.79 K, 333.33 K, 352.94 K, 375.00 K, and 400 K, respectively, with the Langevin middle integrator^65^ and a friction coefficient of 1 ps*^−^*^1^. Each replica lasted for 200 million steps with a timestep of 10 fs. Exchanges between neighboring replicas were attempted every 1000 steps. More details about the replica exchange simulations are attached in the Supporting Information *Section: Implementation of the temperature replica exchange algorithm*. We discarded the first 100 million steps as equilibration and used the remaining data for analysis.

We carried out slab simulations to evaluate the stability of condensates formed by HP1*α* and HP1*β* dimers. Initial configurations of these simulations were prepared as follows. First, we randomly placed 100 copies of protein dimers into a cubic box of length 75 nm. Then we performed 5-million-step constant pressure and constant temperature (NPT) simulations at one bar and 150 K to compress the system with a timestep of 10 fs. Pressure and temperature control were achieved by coupling the Monte Carlo barostat with the Langevin middle integrator.^65^ The length of the compressed cubic box was about 25 nm. Then we fixed the compressed configuration and extended the box size to 25 *×* 25 *×* 400 nm. The rectangular geometry leads to the creation of a dense-dilute interface along the *z*-axis. Starting from this initial configuration, we gradually raised the temperature from 150 K to a target value in the first 0.1 million steps. We then performed 200-million-step production simulations at constant volume and constant temperature using the Nośe-Hoover integrator^65^ with a collision frequency of 1 ps*^−^*^1^ and a timestep of 5 fs.

Following similar protocols outlined above, we performed slab simulations for disordered regions of protein DDX4 and FUS with the HPS model using parameters derived from the Urry hydrophobicity scale.^66^ Detailed amino acid sequences of the two proteins are provided in the Supporting Information. For each protein, we first obtained an equilibrium configuration from a 0.1-million-step constant temperature simulation initialized with a straight C*_α_* chain. We placed 100 replicas of the equilibrium configurations into a cubic box of length 75 nm. Upon compression by a 5-million-step NPT compression at 1 bar and 150 K with a timestep of 10 fs, the system reaches a cubic box with a size of about 15 nm. We then performed slab simulations with an elongated box of size 15 *×* 15 *×* 280 nm^3^ and a 10 fs timestep. Nośe-Hoover integrator was again applied with a collision frequency of 1 ps*^−^*^1^ to maintain the temperature.

### Computing phase diagrams from slab simulations

To determine the concentration of dense and dilute phases from slab simulations, we first identified the largest cluster in a given configuration as the largest connected component of the protein-contact network. Two proteins were defined as in contact if their center-of-mass distance was less than 5 nm. Afterward, we translated the system so that the center of mass of the largest cluster coincides with the box center, which was located at *z* = 0. We recognized the region with *|z| <* 5 nm for HPS simulations and *|z| <* 10 nm for MOFF simulations as the dense phase, while the region with *|z| >* 50 nm as the dilute phase. The concentrations were determined as the average density value in specified regions using the second half of the simulation trajectories. We fitted the concentration values at various temperatures using the following equation to determine the critical temperature

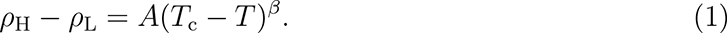

*ρ*_H_ and *ρ*_L_ are the densities at the concentrated and dilute phases. Parameter *β* = 0.325 is the universal critical exponent of 3-dimensional space phase transition. *T*_c_ is the critical temperature and *A* is the coefficient.

## Results

### Flexible biomolecular simulations with diverse force fields

OpenABC implements several existing force fields for coarse-grained (CG) modeling of protein and DNA molecules (Figure 1). These force fields represent each amino acid and nucleotide with one bead. The hydropathy scale (HPS) models for proteins define interactions between different pairs of amino acids based on various hydrophobicity scales.^20,28^ Recent studies attempted to improve the accuracy of HPS models with systematic optimizations of the hydrophobicity scale to match experimental observations of IDP monomers.^34,36^ They have been used to study the phase behaviors of numerous proteins, ^67–69^ revealing the contribution of charge distribution patterns, cation-*π* interactions, and the balance between hydrophobic and electrostatic interactions^68,70^ to condensate stability.

**Figure 1:**
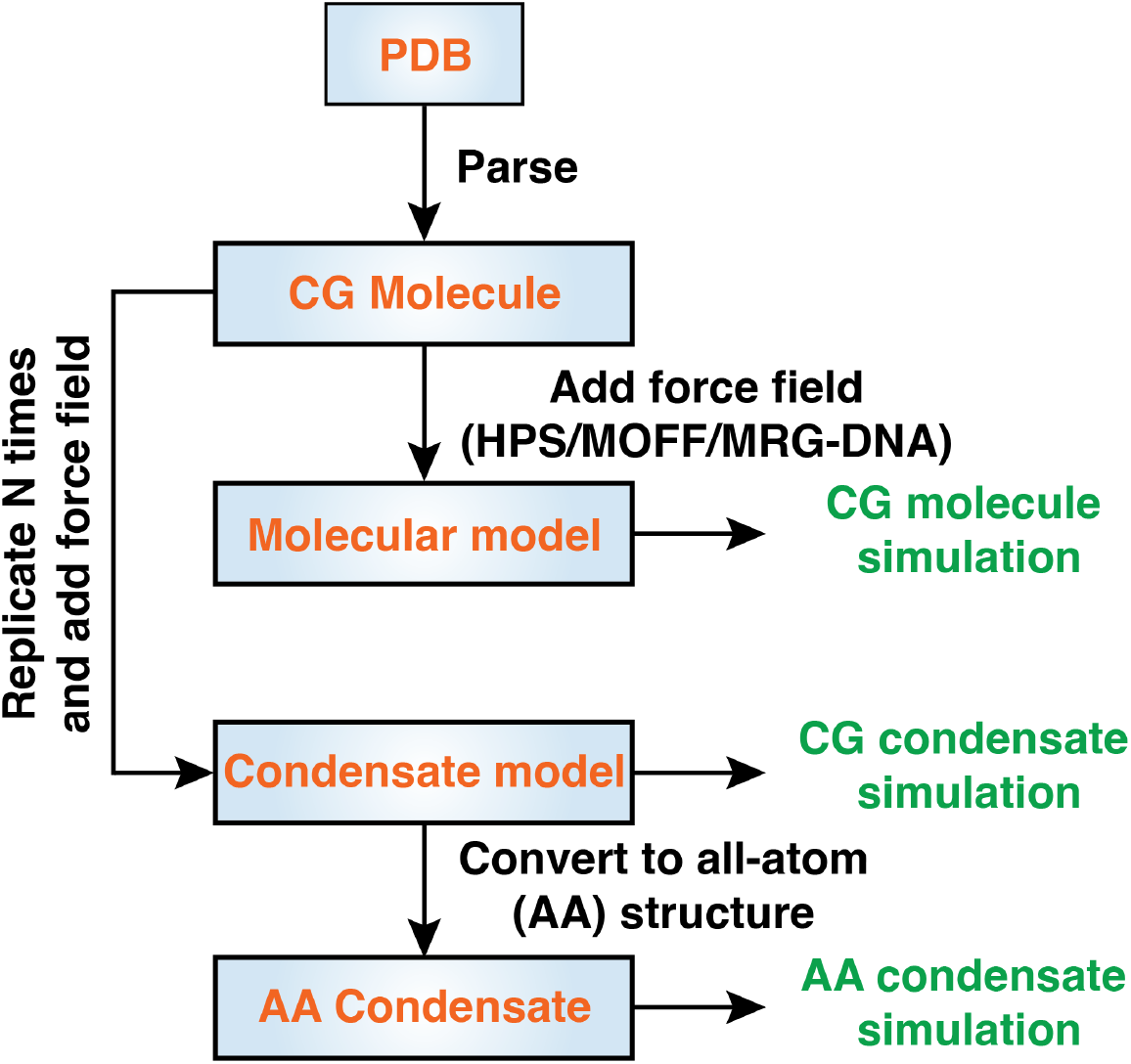
OpenABC facilitates coarse-grained and atomistic simulations of biomolecular condensates with multiple force fields. The diagram illustrates the workflow and various functionalities of OpenABC. To set up condensate simulations, the users must provide a configuration file in the PDB format for the molecule of interest. Open-ABC parses topological and structural information from the PDB file to build a molecule object. Specifying force field options allows direct simulations of individual molecules. On the other hand, the molecule object can be replicated for condensate simulations. In addition, OpenABC allows the conversion of CG configurations to atomistic structures for simulations with all-atom force fields.

We also include MOFF in our OpenABC implementation. MOFF is a protein force field that was parameterized with the maximum entropy algorithm^71,72^ and the protein folding energy landscape theory^73^ to provide consistent descriptions of both folded and disordered proteins.^60,74–76^ It was shown to reproduce the radius of gyration for a collection of proteins, including both ordered and disordered proteins.^60,77^ The balanced interactions among amino acids have proven beneficial for describing the complex contacts among phase-separating proteins, including those with both ordered and disordered domains.^47,60,76^

In addition to protein models, we implemented the molecular renormalization group coarse-graining (MRG-CG) DNA model. This model was parameterized with results from all-atom simulations to reproduce the salt-dependent DNA persistence length.^61^ While MRG-CG DNA was initially introduced for simulations with explicit ions, we adopted it for implicit ion modeling with the Debye-Hückel approximation for electrostatic interactions. We rescaled the strength of bonded interactions to ensure the accuracy of the implicit-ion model in reproducing DNA persistence length at the physiological salt concentration.^47^ It is also worth mentioning that another widely-used CG DNA model, 3SPN.2C,^78^ has already been implemented in OpenMM^79^ and can be readily used.

We modeled protein-DNA interactions with both excluded volume effects and electrostatic interactions. Previous studies have shown that these treatments successfully quantify the stability of various protein-DNA complexes, ^37,41,79–84^ revealing the complex phase behavior of protein-DNA condensates.^47,85^ Detailed expressions of the energy functions for all the force fields are provided in the Supporting Information *Section: Force Field Definitions*, with the parameters provided in Tables S1-S6.

### Simplified Setup of Condensate Simulations

OpenABC leverages the MD simulation engine, OpenMM,^62^ to offer simulation setup with Python scripting, thus dramatically simplifying the workflow. The software treats each molecule as an object and appends such objects into a container-like class. This class allows the incorporation of various force field options and integration schemes for MD simulations.

An illustration of the typical workflow for condensate simulations is provided in Figure 1. OpenABC first parses a configuration file in the PDB format supplied by users to create a molecule object. The object contains topological and structural information extracted from the input file. Upon introducing interactions defined in various force fields, the molecule object can be used to simulate individual biomolecules. On the other hand, the molecule object can also be replicated *N* times for condensate simulations consisting of *N* molecules. As demonstrated in an example code in Figure 2, setting up an entire MD simulation of a protein condensate with default parameters only requires about 20 lines of code.

**Figure 2:**
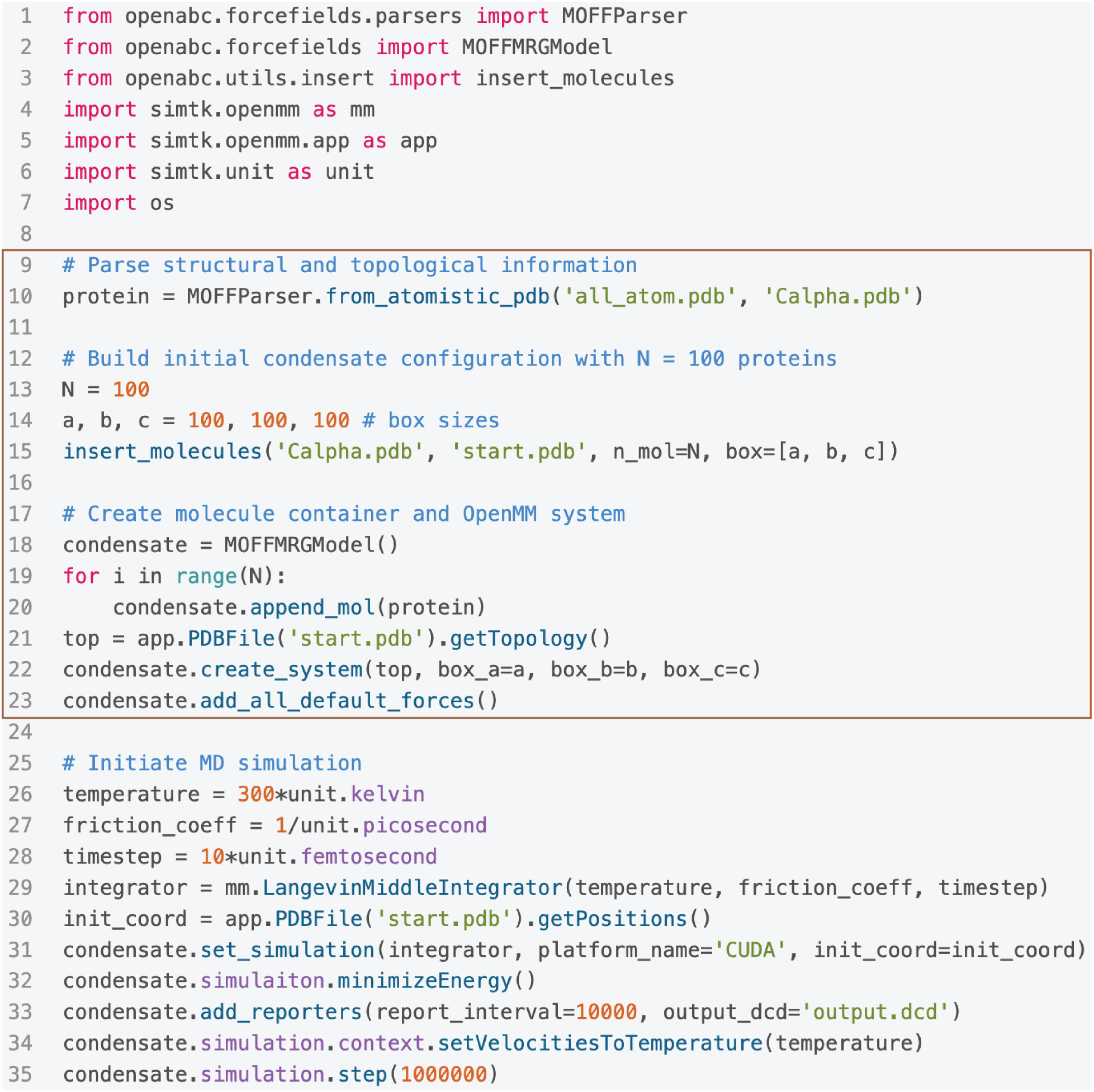
OpenABC simplifies simulation setup with Python scripting. The example code includes all the steps necessary for setting up and performing MD simulations of a protein condensate with MOFF and default settings in a cubic box of length 100 nm. The ten lines included in the highlight box correspond to the creation of the condensate system by parsing topological information from an initial PDB file, building a configuration file by inserting molecules into a box and incorporating the molecular objects, *protein*, into a container class, *condensate*, with appropriate force fields. The rest of the code includes standard simulation setups generic to OpenMM. We chose the Langevin middle integrator to perform simulations at 300 K with a friction coefficient of 1 ps*^−^*^1^ and a timestep of 10 fs.

To enhance conformational sampling of individual molecules and condensates, we provide an implementation of the temperature replica exchange algorithm^63^ with PyTorch^86^ as part of the package (see Supporting Information *Section: Implementation of the temperature replica exchange algorithm* for details). Furthermore, we introduce utility functions to reconstruct atomistic structures from coarse-grained protein configurations with only *α* carbons. This functionality relies on the software “reconstruct atomic model from reduced representation (REMO)”^87^ and can facilitate downstream all-atom simulations. More tutorials in Jupyter Notebook format are available online at the OpenABC GitHub repository.

### Efficient simulations with GPU-enabled MD engine

A significant advantage of integrating with OpenMM comes from its native support of GPU acceleration. Simulating coarse-grained condensates on GPUs can be particularly beneficial due to the inhomogeneous distribution of particles arising from implicit solvation. ^79^ CPU parallelization, which often relies on the spatially-based, domain decomposition strategy, is often less effective because the inhomogeneity in particle density between the condensate and solvent phases produces an imbalanced workload between CPUs.

To demonstrate the efficiency of GPU-enabled simulations, we studied five independent condensate systems. The first four systems consist of *N*_1_ HP1*α* dimers and *N*_2_ 200-bp-long dsDNA randomly distributed in a cubic box of length 200 nm with periodic boundary conditions. In the fifth system, 100 HP1*α* dimers in a compact configuration were placed at the center of an elongated box of size 25 *×* 25 *×* 400 nm. This rectangular setup is typical for the so-called slab simulations to produce a dilute and dense interface along the *z*-axis for computing co-existence curves and phase diagrams. ^20,88,89^ MOFF and MRG-CG force fields were used to describe the interactions among coarse-grained particles. We simulated each system for one million timesteps using the Langevin middle integrator^65^ to control the temperature at 300 K, with a friction coefficient of 1 ps*^−^*^1^ and a time step of 10 fs. For comparison, we simulated the same systems with a closely related integrator using GROMACS, a leading MD engine with state-of-the-art performance on CPUs. ^54,57^ More simulation details are provided in the Supporting Information *Section: Benchmarking the performance of condensate simulations*.

As shown in Figure 3, OpenMM single GPU performance matches GROMACS with hundreds of CPUs in the first four systems. While GROMACS achieved nearly linear scaling for the first four systems, introducing more CPUs did not lead to any significant speedup in the last system with a dense-dilute interface. As mentioned above, the presence of vacuum regions in slab simulations hinders the efficacy of domain decomposition. On the other hand, OpenMM is less sensitive to the simulation setup and retains superior performance.

**Figure 3:**
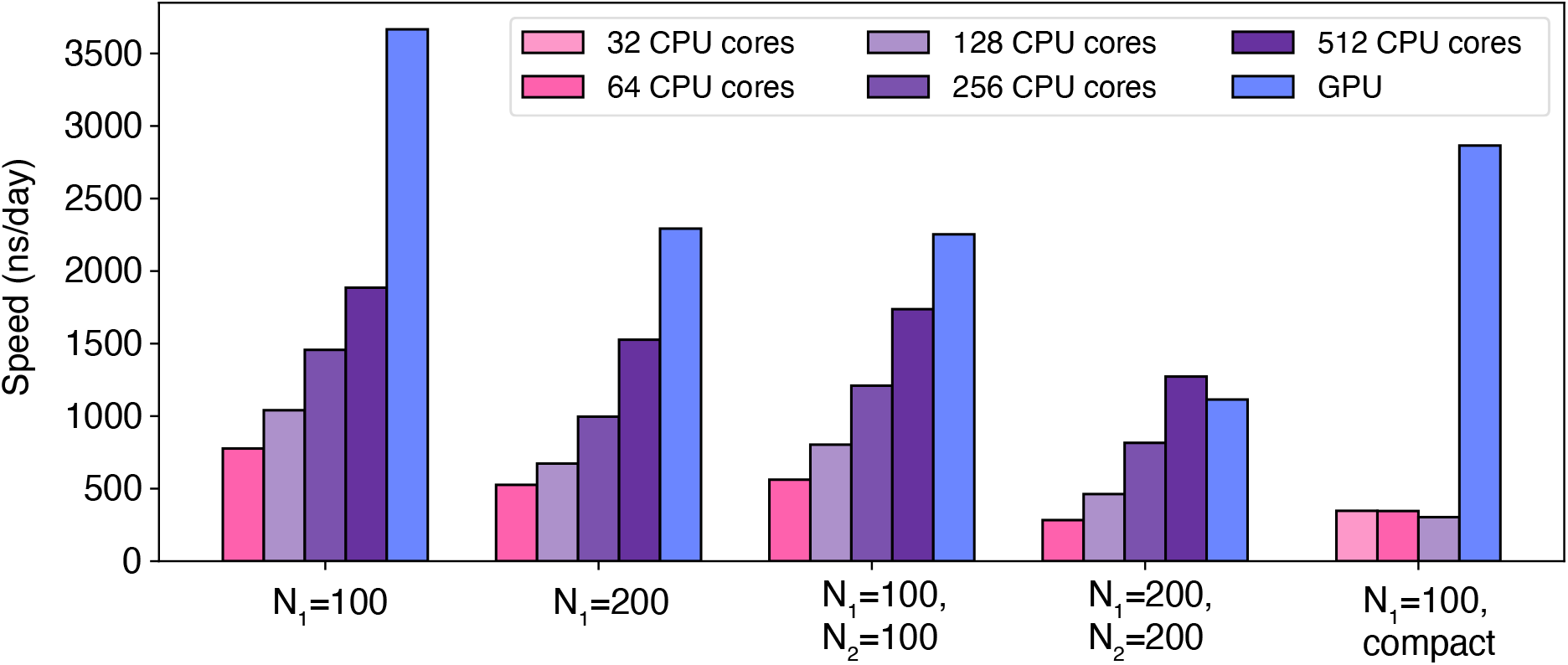
OpenABC integrates with OpenMM for GPU-accelerated MD simulations. The five data sets compare the performance of CPU simulations using GROMACS with single GPU simulations using OpenMM. The different colors indicate the number of CPUs in GROMACS simulations, as shown in the legends. The simulation systems consist of *N*_1_ HP1*α* dimers and *N*_2_ 200-bp-long dsDNA (*N*_2_ = 0 if not specified). The first four systems adopt homogeneous density distributions in cubic boxes, while the last exhibits a dense-dilute interface. The benchmarks were performed with Intel Xeon Gold 6248 CPUs or Intel Xeon Platinum 8260 GPUs.

### Application: Validating force field implementation in OpenABC

Before applying the software for extensive simulations, we validated our implementation of various force fields. We generated ten configurations for an HP1*α* dimer with MOFF and GROMACS through an NVT simulation. As shown in Table S7, the potential energies evaluated using MOFF from OpenABC match those reported by GROMACS. Similar comparisons using a protein-DNA complex produce nearly identical energy values as well. The protein-DNA complex is formed by an HP1*α* dimer with a 200-bp-long dsDNA, and MOFF and MRG-CG DNA were used to quantify their interactions. The minor differences between OpenMM and GROMACS energies are mainly caused by using tabulated functions for nonbonded interactions in GROMACS.

We further evaluated the potential energies defined by the HPS model on ten configurations of a disordered protein, DDX4, using both OpenMM and HOOMD-Blue.^90^ As shown in Table S9, the two sets of energies match exactly, supporting the correctness of our force field implementation.

In addition to energy comparisons, we further examined the conformational ensembles of HP1*α* and HP1*β* dimers using MOFF with temperature replica exchange simulations. ^63^ Consistent with our previous study,^60^ the force field succeeds in resolving the difference in their conformational distribution between the two homologs (Figure 4). The radius of gyration for the two dimers at 300 K is 3.33 *±* 0.19 nm, and 4.27 *±* 0.09 nm, respectively. These values match the previously reported values computed using GROMACS quantitatively, reproducing experimental trends. Therefore, OpenABC produces consistent results with other software despite differences in integration schemes.

**Figure 4:**
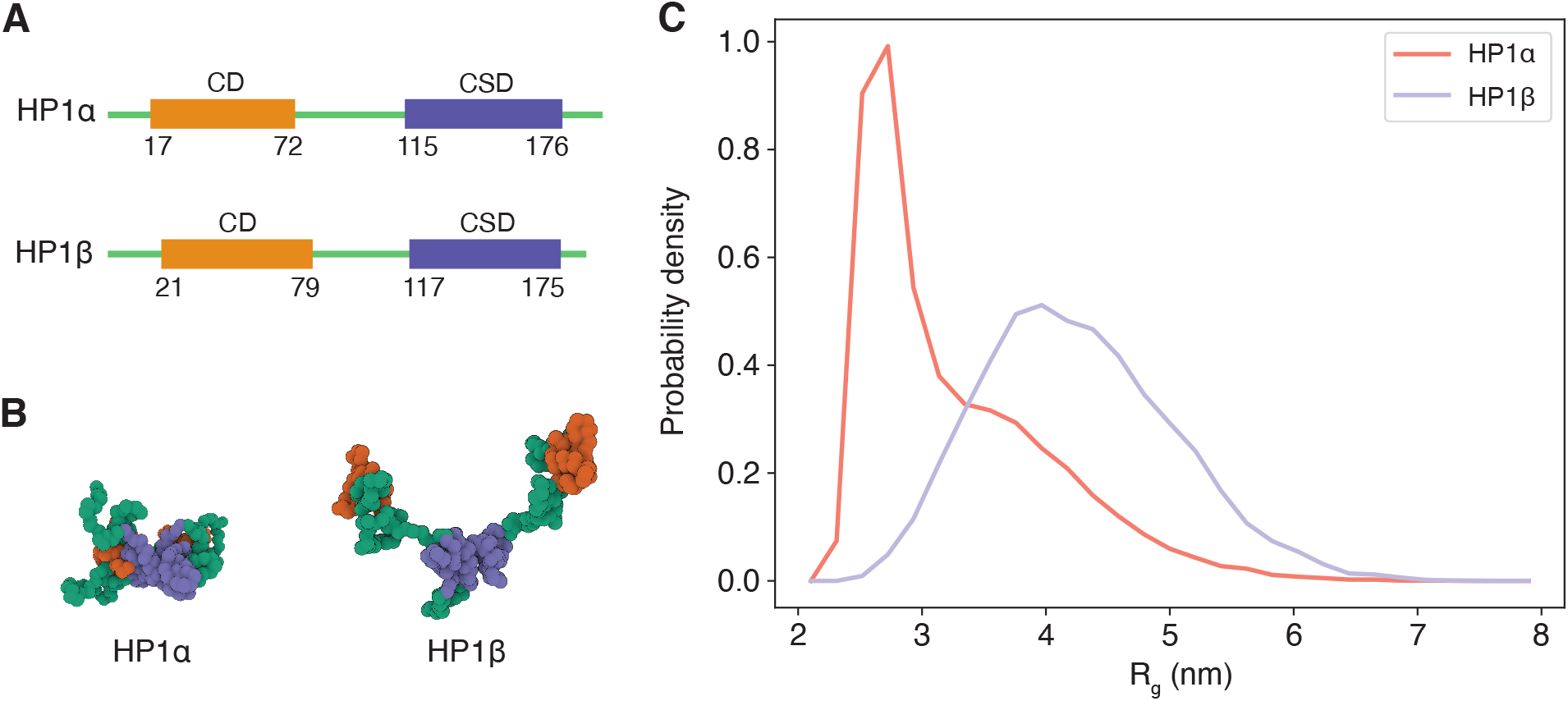
OpenABC produces consistent results with a previous studying, resolving the structural differences between two HP1 homologs. (A) Secondary structures of HP1*α* and HP1*β* along sequences. (B) Representative structures for HP1*α* and HP1*β* dimer rendered with Mol* Viewer.^91^ The radius of gyration (*R_g_*) for the two structures are 2.77 and 4.44 nm, respectively. We colored the chromodomain (CD) in orange, the chromoshadow domain (CSD) in blue, and the rest in green. (C) Probability density distributions of *R_g_* for HP1*α* (red) and HP1*β* dimer (blue).

### Application: Coarse-grained simulation of protein condensates

As additional evaluations of force field implementation and to demonstrate the usefulness of OpenABC, we performed slab simulations to determine the phase diagram of four proteins. The simulations for HP1*α* and HP1*β* were performed with MOFF, while those for FUS LC and DDX4 were modeled with the HPS model using the shifted Urry hydrophobicity scale.^28^ The resulting phase diagrams are shown in Figure 5, with the numerical values listed in Tables S10 and S11. The density profiles at different temperatures are shown in Figures S2 and S3. We fitted the computed phase diagrams with an analytical expression to determine the critical temperature *T_c_* (see Methods). The critical temperatures are 306.30 K for HP1*α* and 245.99 K for HP1*β*, consistent with previous results obtained with GROMACS simulations.^60^ Similarly, the critical temperatures for DDX4 and FUS LC are 324.21 K and 340.04 K, respectively, matching values reported in a previous study that used the software HOOMD-Blue for simulations. ^28^ Thus, OpenABC produces statistically indistinguishable results on the phase behavior of protein condensates as in previous studies.

**Figure 5:**
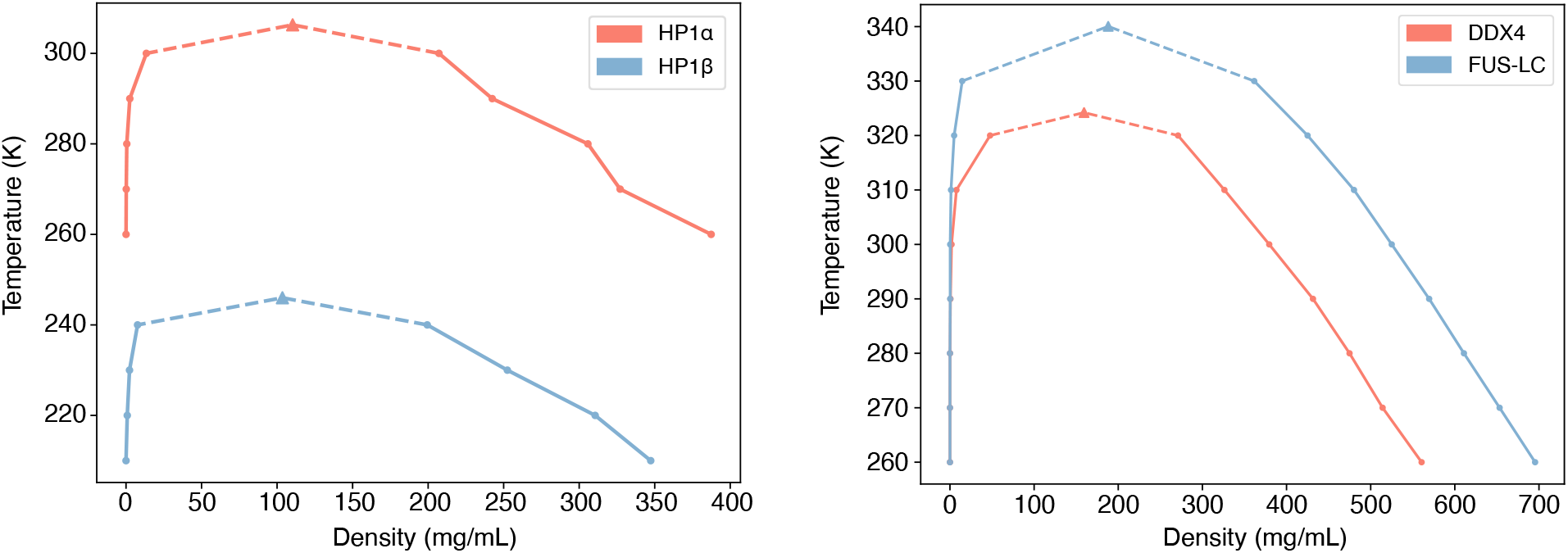
OpenABC produces phase diagrams that match previous results. (A) Phase diagrams for HP1*α* (red) and HP1*β* (blue) dimer condensates computed with MOFF. (C) Phase diagrams for DDX4 (red) and FUS LC (blue) computed with the HPS model parameterized using the Urry hydrophobicity scale. The dots in both plots denote the density values determined from slab simulations, and the triangles represent the critical point obtained from numerical fitting.

### Application: Atomistic simulation of protein condensates

While residue-level coarse-grained models are helpful for long timescale simulations, their limited resolution may prove insufficient for characterizing specific condensate properties, including the solvation environment,^48^ counter ion distribution,^49^ and protein-ligand interactions.^50^ Therefore, we implemented functionalities in OpenABC to convert equilibrated CG configurations to atomistic structures for downstream all-atom simulations when desired.

As proof of principle, we converted the final snapshot from the slab simulation of HP1*α* dimer at 260 K to an atomistic configuration (Figure 6). This conversion leverages the software REMO^87^ to build atomistic details starting from the *α*-Carbon position of each amino acid. We further solvated the atomistic structure with water molecules and counter ions. After energy minimization, we carried out an all-atom MD simulation using GROMACS with the CHARMM36m force field^92^ and the CHARMM-modified TIP3P water model.^93^ More details about simulation preparation can be found in the Supporting Information *Section: Building and relaxing atomistic structures from coarse-grained configurations*. As shown in Figure 6, the system relaxes with a continuously decreasing potential energy in the first 20 ns and remains stable afterward.

**Figure 6:**
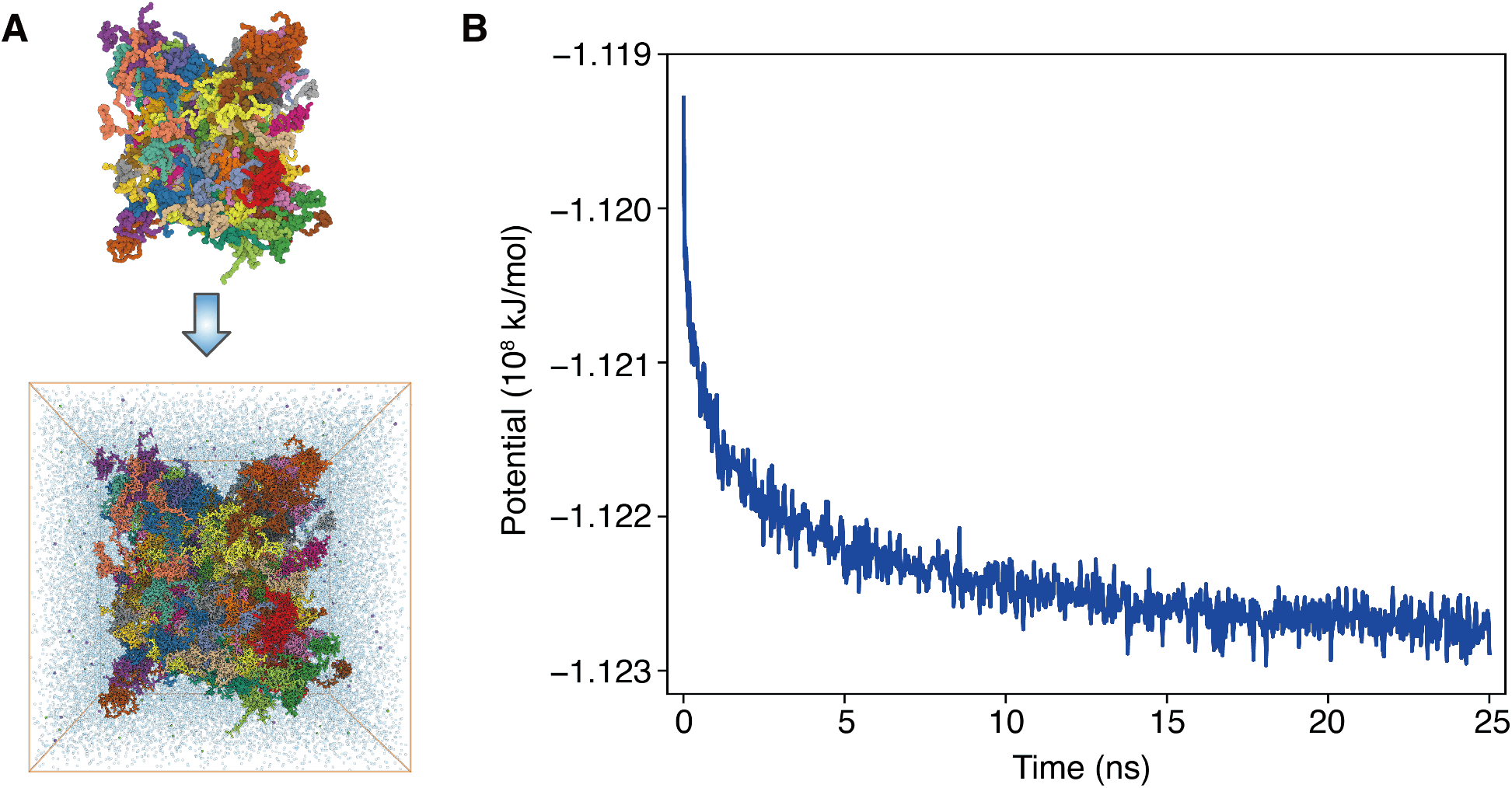
OpenABC facilitates all-atom simulations by producing equilibrated initial atomistic configurations. (A) Illustration of the conversion from a coarse-grained configuration (top) to a fully atomistic model with explicit solvent molecules (bottom). Only 2% of water molecules and counter ions of the atomistic model are shown for clarity. The system consists of 100 HP1*α* dimers, and different molecules are shown in one of 25 colors. Both figures are rendered with Mol* Viewer.^91^ (B) The atomistic potential energy evaluated using the CHARMM force field is shown as a function of simulation time.

## Conclusions

We introduced a software package, OpenABC, to facilitate coarse-grained and all-atom simulations of biomolecular condensates. The package implements several of the leading coarsegrained force fields for protein and DNA molecules into OpenMM, enabling GPU-accelerated simulations with performances rivaling GROMACS simulations with hundreds of CPUs. New force fields can be quickly introduced within the framework, and we plan to incorporate RNA models into the package as the next step. Comprehensive tutorials are provided to familiarize the users with the various functionalities offered by OpenABC. We anticipate the intuitive Python interface of OpenABC to reduce the entry barriers and promote coarse-grained modeling for its adoption by a broader community.

## Acknowledgement

This work was supported by the National Institutes of Health (Grant R35GM133580) and the National Science Foundation (Grant MCB-2042362). We appreciate Azamat Rizuan’s help verifying the implementation of HPS models.

## Competing interests

The authors declare that they have no competing interests.

## Data and materials availability

OpenABC source code is available at https://github.com/ZhangGroup-MITChemistry/OpenABC

## Notes

### Competing Interest Statement

The authors have declared no competing interest.

